# Assessment of the acute effects of 2C-B vs psilocybin on subjective experience, mood and cognition

**DOI:** 10.1101/2023.02.16.528808

**Authors:** Pablo Mallaroni, Natasha L. Mason, Johannes T. Reckweg, Riccardo Paci, Sabrina Ritscher, Stefan W. Toennes, Eef L. Theunissen, Kim P.C. Kuypers, Johannes G. Ramaekers

## Abstract

2,5-dimethoxy-4-bromophenethylamine (2C-B) is a hallucinogenic phenethylamine derived from mescaline. Observational and preclinical data have suggested it to be capable of producing both subjective and emotional effects on par with other classical psychedelics and entactogens. Whereas it is the most prevalently used novel serotonergic hallucinogen to date, it’s acute effects and distinctions from classical progenitors have yet to be characterised in a controlled study. We assessed for the first time the immediate acute subjective, cognitive, and cardiovascular effects of 2C-B (20 mg) in comparison to psilocybin (15mg) and placebo in a within-subjects, double-blind, placebo-controlled study of 22 healthy psychedelic-experienced participants. 2C-B elicited alterations of waking consciousness of a psychedelic nature, with dysphoria, subjective impairment, auditory alterations, and affective elements of ego dissolution largest under psilocybin. Participants demonstrated equivalent psychomotor slowing and spatial memory impairments under either compound compared to placebo, as indexed by the Digit Symbol Substitution Test (DSST), Tower of London (TOL) and Spatial Memory Task (SMT). Neither compound produced empathogenic effects on the Multifaceted Empathy Test (MET). 2C-B induced transient pressor effects to a similar degree as psilocybin. The duration of self-reported effects of 2C-B was shorter than that of psilocybin, largely resolving within 6 hours. Present findings support the categorisation of 2C-B as a subjectively “lighter” psychedelic. Tailored dose-effect studies are needed to discern the pharmacokinetic dependency of 2C-B’s experiential overlaps.

## Introduction

Classical psychedelics are a diverse set of psychoactive compounds characterised by their capacity to elicit profound alterations in waking consciousness, comprising changes to mood, cognition, and self-referential awareness^1^. Promising evidence that compounds such as psilocybin are safe and efficacious treatments for neuropsychiatric disorders such as major depressive disorder (MDD) has fostered a renewed interest in their effects^2-4^.

Like other psychedelics (e.g. *d*-lysergic acid - LSD, mescaline, and *N,N*-dimethyltryptamine - DMT), the effects of psilocin (the active metabolite of psilocybin) are understood to be primarily mediated via partial serotonin (5-HT) receptor agonism, predominantly the 5-HT_2A_ receptor as well as 5-HT_1A_ and 5-HT _2C_ receptors ^5,6^. Extensive human behavioural studies have characterised the acute subjective, cognitive, and autonomic effects of psilocybin ^7-13^, making it a valuable candidate for clinical use.

While significant emphasis has been placed on the central role of 5-HT_2A_ receptor agonism in mediating alterations to waking consciousness ^14^, classical psychedelics are suggested to hold subtly distinct pharmacodynamics. Indoleaklymaines such as the ergoline LSD or the tryptamines psilocybin and DMT exhibit varying receptor binding selectivity and potency particularly in comparison to phenethylamines such as mescaline, likely as a function of their structural (as)similarity ^15,16^. Evidence has begun to emerge suggesting experiential differences may not only be explained by functional selectivity at the 5-HT_2A_ receptor but in contrast, orchestrated by an entourage of receptor subclasses^17,18^

With the advent of modern biochemistry, novel compounds distinct from their classical progenitors have readily appeared. Of these is the psychedelic entactogen 2,5-dimethoxy-4-bromophenethylamine (2C-B), a first-generation synthetic analogue of mescaline isolated and publicised by Alexander Shulgin in 1974^19^. Belonging to the 2C-series of 2,4,5 trisubstituted phenethylamines, members share core methoxy functional groups at carbons 2 and 5, but differ in ligands at the fourth position on the phenyl ring, e.g., bromine in 2C-B or propyl in 2C-P ^20^. Manifold increases in receptor efficacy and potency with respect to mescaline have been observed following such alterations in ligand positioning ^21^. 2C-B shows high 5-HT2A/5-HT_2C_ receptor selectivity and has been demonstrated to reliably elicit head twitch responses in rodents, a behavioural proxy of 5-HT_2A_ agonism ^22-25^. In recent years, research has indicated secondary affinity at 5-HT_2B_,5-HT_1A_, dopamine D_1/2/3_, adrenergic α_1/2_, histamine H_1_ and TAAR_1_ receptors^21^. Notably, 2C-B shows parallels with 3,4-methylenedioxymethamphetamine (MDMA) due to (albeit weaker) reuptake inhibition of the serotonin reuptake transporter (SERT) and, to a lesser extent, norepinephrine (NET) and dopamine reuptake transporters (DAT) ^26,27^, likely supplemented by monoamine oxidase (MAO_A/B_) inhibition ^28,29^.

To date, 2C-B is the most frequently used novel psychedelic among recreational drug users ^30,31^ and serves as a structural reference point for the development of novel compounds ^32^. Despite its prevalence, little is known regarding its effect profile in humans, making it difficult to ascertain any potential risks or clinical merits. A body of survey data and anecdotal reports highlight mild psychedelic effects typical of serotonergic hallucinogens, paired with feelings of euphoria and emotional openness akin to the entactogen MDMA^33,34^. Whereas naturalistic observational studies have begun to corroborate these reports and expanded on the relative safety of 2C-B ^35,36^, a complete neuropsychopharmacological assessment of its acute effects under double-blind conditions has yet to be performed.

The present study therefore sought to describe and compare the subjective, cognitive, and cardiovascular effects of 2C-B (20 mg) vs psilocybin (15 mg) psilocybin in healthy volunteers. Comparative studies employing a within-subjective design offer the opportunity to directly assess commonalities between substances across a range of standardised measures. Moreover, by including different compounds of comparable phenomenology and effect duration, it may also be possible to minimise expectancy effects clouding subjective responses to psychedelics ^37^. Considering prior findings of entactogenic qualities, we hypothesised 2C-B would produce distinct subjective emotional effects compared to psilocybin.

## Materials and methods

The study employed a double-blind, placebo-controlled, crossover design with three acute experimental sessions to investigate responses to placebo, 2C-B, and psilocybin. Participants were randomly allocated to intervention orders following Latin-square counterbalancing. Each acute session was separated by a 14-day washout. The study (trial register NL8813) was conducted according to the Declaration of Helsinki (1964) and amended in Fortaleza (Brazil, October 2013) and in accordance with the Medical Research Involving Human Subjects Act (WMO) and was approved (NL73539.068.20) by the Academic Hospital and University’s Medical Ethics committee of Maastricht University.

### Participants

Twenty-two healthy participants (11 female) aged 19-35 years (mean ± SD: 25 ± 4 years) were recruited by word of mouth and advertisement shared via Maastricht University social media platforms.

The inclusion criteria were: 18-40 years of age; previous experience with a psychedelic drug but not within the past 3 months; body mass index between 18 and 28 kg/m2; free from medication (any drug prescribed for a medical indication); good physical health, including absence of major medical, endocrine, and neurological conditions; and written informed consent. The exclusion criteria included history of drug abuse or addiction, pregnancy, or lactation, current or history of psychiatric disorders, absence of reliable contraceptives, previous experience of serious side effects to psychedelics, and MRI contraindications. Before inclusion, participants were screened and examined by an independent study physician, who checked for general health, conducted a resting ECG, and took blood and urine samples in which haematology, clinical chemistry and urine analyses were conducted.

All participants were fully informed of all procedures, possible adverse reactions, legal rights, and responsibilities, expected benefits, and their right to voluntary termination without consequences. All participants provided their informed consent, in writing, prior to their inclusion in the study and are to be remunerated for their participation. An overview of the recruitment process, CONSORT flowchart and complete demographic data can be found in the supplementary materials.

### Study drugs

Each intervention was administered orally, in a closed cup either containing placebo (inactive; bittering agent) or bittering agent and psilocybin/2C-B (powder). Blinding integrity was assessed retrospectively at the end of each acute experimental visit. A permit for obtaining, storing, and administering 2C-B and psilocybin was obtained from the Dutch Drug Enforcement Administration. The choice of dose and drug was determined by the requirement to generate substantial, yet tolerable, altered state of consciousness in the absence of existing 5-HT_2A_ receptor occupancy data for 2C-B. Prior work has indicated the tolerability, safety and comparable onset times of moderate dose of 20 mg 2C-B ^36,38^ and 15mg psilocybin ^9,39^.

### Study procedures

Upon enrolment, and prior to the start of testing cycles, participants were invited to a preparatory training visit where they met and built rapport with the experimenters and were trained on all computerised tasks and procedures.

Each acute experimental session lasted approximately 7 hours. On the morning of the dosing day, participants were instructed to have a light breakfast at home. Participants were reminded to refrain from drug use, including psychedelic drugs (≥ 3 months), alcohol (≥24 hours), and all other drugs of abuse (≥7 days) before each experimental visit. Additionally, participants were asked to refrain from caffeine and nicotine use on each test day. All psychoactive drug use was prohibited throughout the duration of the study (7 weeks). On arrival of a test day, absence of drug and alcohol use was assessed via a urine drug screen and a breath alcohol screen. A pregnancy test was given to females. If all tests were found to be negative, a venous catheter was placed, and participants were allowed to proceed with administration. The acute experimental sessions were conducted in a quiet psychopharmacology lab. In between testing, the participants could interact with the investigator, rest, read, or listen to music via headphones. Participants stayed under supervision until the testing day was complete and the experimenters deemed they were fit to go home.

### Subjective effects

Detailed descriptions of all inventories are provided in the supplementary materials.

Subjective effects were assessed repeatedly using visual analogue scales (VAS) at baseline (0h), +0.5, +1, +1.5, +2, +3, +4, +5, +6 hours after administration. These scales included “*any drug effect*,” “*good drug effect*”, “*bad drug effec*t”, “*drug liking*,” “*drug high*”,” “*happy*”,” *concentration*”, *“creative*”, *“productive*”, ”*sociable*” marked from “not at all” (0) to “extremely” (100) and *“sense of time*” marked from “slow” (0) to “fast” (100) ^40,41^.

Hourly measures (0, +1, + 2, +3, +4, +5, +6h) were taken for a broader characterisation of present-state effects. Changes in mood were assessed using the 65-item Profile of Mood States (POMS) scale ^42^. Levels of dissociative symptomatology were examined using the 19-item Clinician-Administered Dissociative States (CADSS) Scale ^43^. Acute psychedelic effects were characterised using the 13-item Bowdle Visual Analogue (BVAS) Scale ^44^. Lastly, current and general levels of drug liking and wanting were assessed using the 4-item Sensitivity to Drug Reinforcement Questionnaire (SDRQ)^45^.

Furthermore, participants were asked to complete retrospective measures at the end of each test day. Alterations in waking consciousness were assessed using the 94-item 5-Dimensional Altered States of Consciousness Rating Scale (5D-ASC) ^46^ and the Hallucinogen Rating Scale (HRS) ^47^. In addition, levels of subjective ego-dissolution as defined by the 8-item Ego Dissolution inventory (EDI) ^48^ and end of session changes in trait empathy as per the 28-item Interpersonal Reactivity Index (IRI) ^49^ were examined.

### Cognitive tasks

A computer-based task battery was implemented during peak subjective effects to determine the differential impact of psilocybin and 2C-B on domains of cognition: Motor Control Task (MCT, sensorimotor coordination)^50^, Psychomotor Vigilance Task (PVT, sustained attention)^51^, Digit Symbol Substitution Test (DSST, overall cognitive impairment)^52^, Tower of London (executive functioning)^53^, the immediate and delayed (+30 min), Spatial Memory Test (SMT, spatial memory)^54^, the Matching Familiar Figures Test (reflection impulsivity)^55^ and the Multifaceted Empathy Test (MET, cognitive and emotional empathy)^56^. Parallel versions of the DSST, TOL, SMT and MFFT were provided for each dosing day. Participants were refamiliarised with all instructions prior to each task. All tasks and corresponding outcome measures are described in detail in the supplementary methods, with times of administration outlined in the upper panels of Fig.4.

### Cardiovascular effects

Blood pressure and heart rate were assessed hourly (0, +1, + 2, +3, +4, +5, +6h) with an arm blood pressure cuff (M6 Comfort Model HEM-7360-E, Omron-Healthcare, Kyoto Japan). For each timepoint, rate pressure product (systolic blood pressure * heart rate) was calculated as a measure of hemodynamic workload ^57^.

### Pharmacokinetic assessments

Venous blood samples were taken hourly (0, +1, + 2, +3, +4, +5, +6h) to assess levels of serum psilocin and 2C-B (ng/mL) using liquid chromatography-tandem mass spectrometry liquid chromatography-tandem mass spectrometry (LC-MS/MS; Agilent, Waldbronn, Germany, see supplementary materials).

### Statistics

We firstly determined peak changes from baseline for all repeated measurements peak effects (*ΔE*_max_ and/or *ΔE*_min_). Baseline-adjusted E_max_ scores were calculated for all cardiovascular measures to assess clinically significant changes. All scores and task outcomes were incorporated into Linear Mixed Models (LMMs) with drug condition as a main fixed effect and participant as a random intercept to account for the dependency between repeated measures. A first-order autoregressive covariance structure (AR1) was used. If a significant main effect of drug condition was observed, estimated marginal means were compared between drug conditions. Tukey’s method was used to correct for multiple comparisons in all analyses.

Prior to pharmacokinetic analyses, incomplete time courses were interpolated using multiple imputation chain equations (MICE), derived from the mice (version 3.15.0) package^58^, following no detection of deviation from missing completely at random (MCAR) based on Little’s MCAR test. Preliminary serum area under the curve (AUC) and half-lives were extrapolated using the NonCompart (version 0.6.0) package^59^. All pharmacokinetic analyses were performed in a non-compartmental fashion. The duration of acute subjective effects was estimated using the VAS “*any drug effect*” and extrapolating an on/off cut-off of 10% of the maximum individual response.

Statistical models were estimated using the *lme* function of the nlme (version 3.1.162) and the ImerTest (version 3.1.3) library ^60,61^ within the R environment ^62^ (version 4.2.1). For all analyses, the alpha criterion was set at *p <* 0.05. Information pertaining to missing data and study power calculation are outlined in the supplementary materials. All statistics are summarised in Table 1 and Table S1-3, with all pairwise comparisons supplied in Table S4. Measures subject to a main effect of drug condition are reported in text.

**Table 1.**
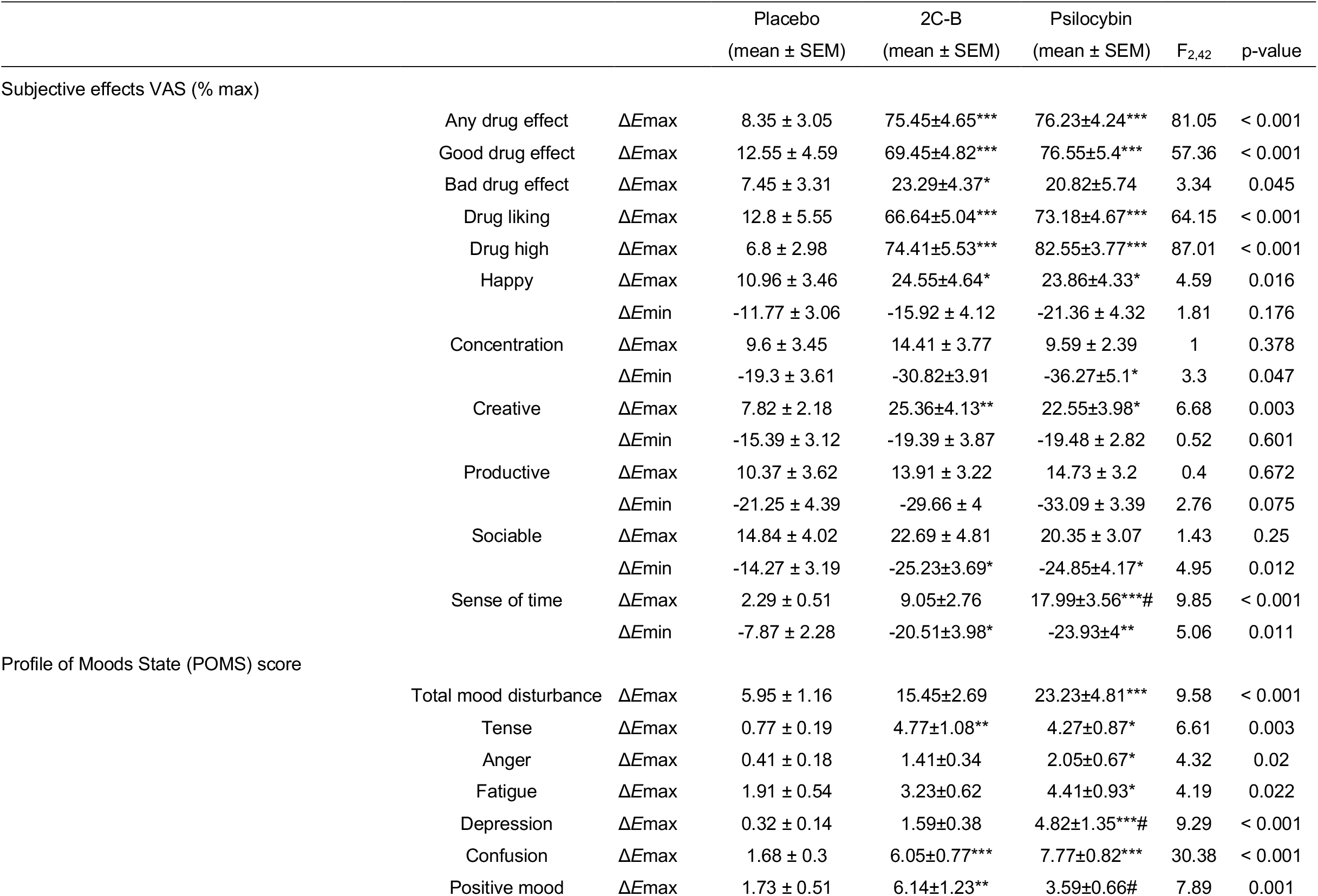
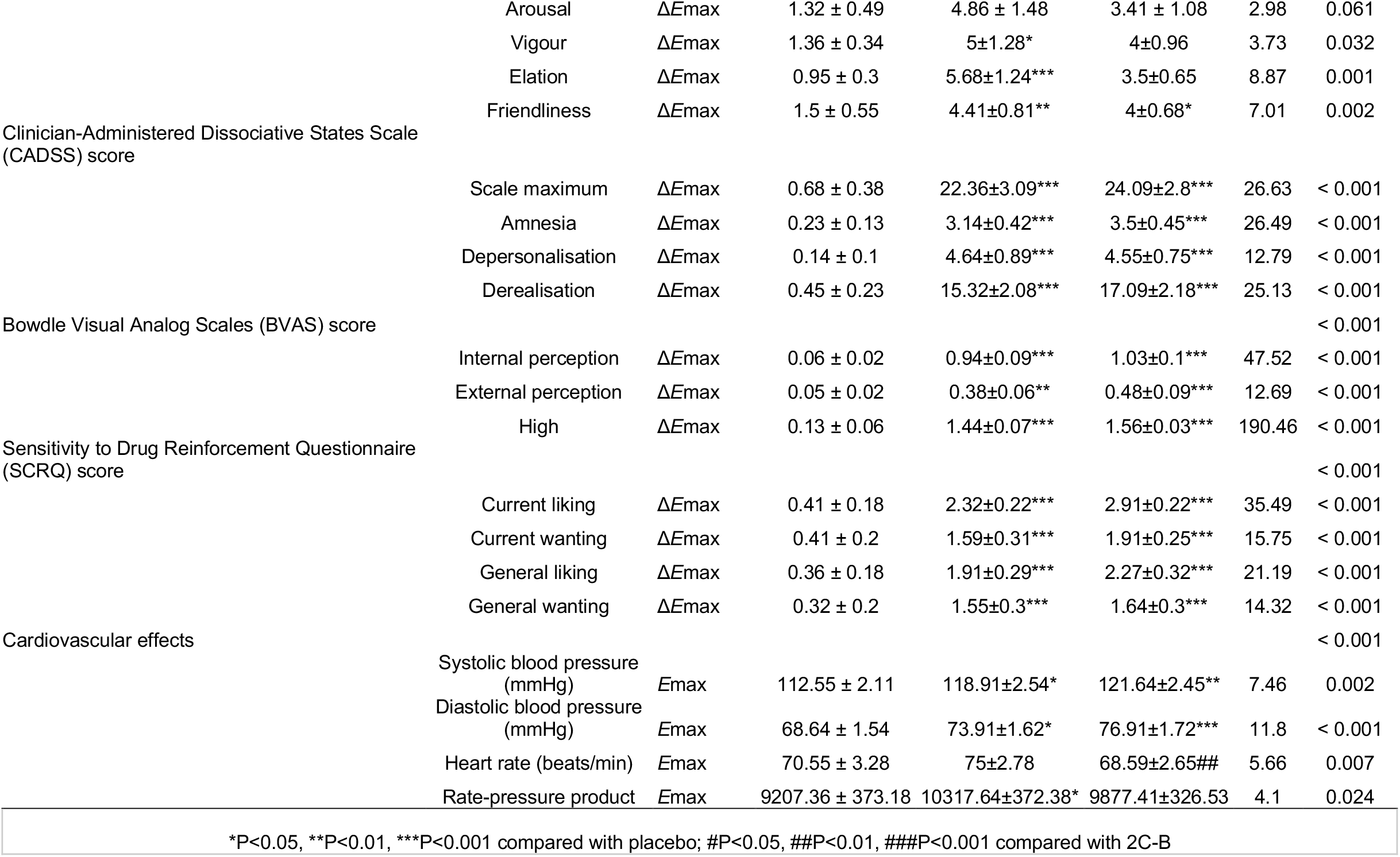
Comparison of the acute effects of 2C-B, psilocybin and placebo

## Results

All 22 participants completed each condition of the study. Both 2C-B and psilocybin were well tolerated, with no serious adverse events arising throughout the course of the study.

### Subjective effects

Subjective effect VAS over time is shown in Fig.1a. Statistics are summarised in Table 1 and Table S4. Overall, 2C-B and psilocybin generated significant yet equivalent increases in scores across most measures of subjective drug intensity (any drug effect, good drug effect, drug liking, drug high) in comparison to placebo. 2C-B produced significant elevations in bad drug effect compared to placebo, but not psilocybin. Subjective effects abated in under 6 hours for 86.3 % of participants under 2C-B vs 63.6 % under psilocybin (Fig.1b, see supplementary for additional details).

**Figure 1.**
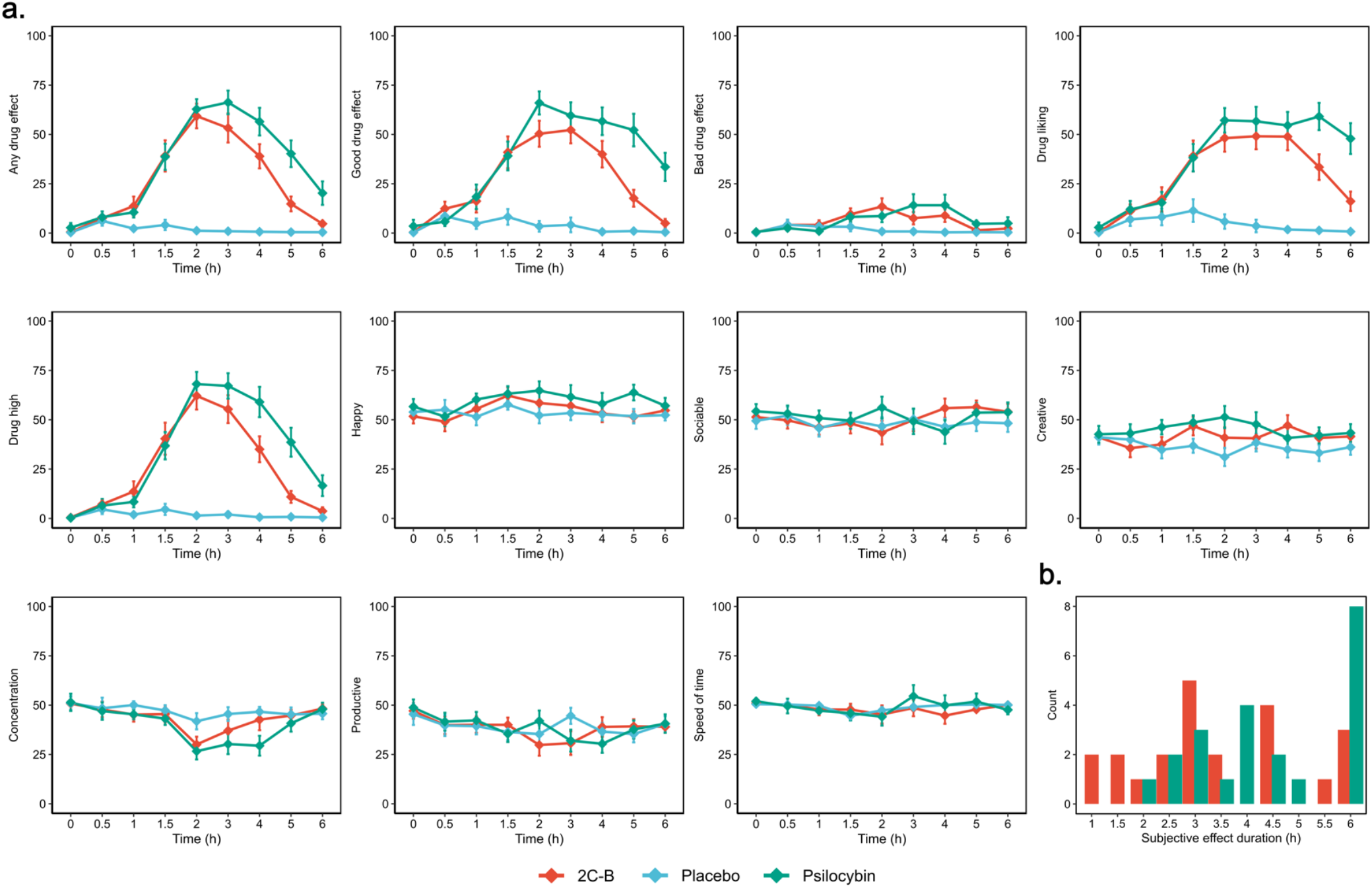
Subjective effects of 2C-B and psilocybin over time on VAS measurements of subjective effects. **(a)** Time courses of each VAS measurement. Data are presented as mean ± SEM (brackets). All values corresponding pairwise statistics are presented in tables 1 and S4. **(b)** Count plot of “Any effect” on/off duration. Absolute frequencies for each bin are depicted for each drug.

VAS pertaining to mood and time perception revealed further similarities. Both drugs significantly increased ratings of happiness and creativity in comparison to placebo with no differences arising between the two. Similarly, 2C-B and psilocybin equally induced greater mean reductions over time for self-reported sociability. Maximal reductions in concentration were significantly greater for psilocybin compared with placebo but not 2C-B. 2C-B and psilocybin similarly induced greater slowing of time perception in comparison to placebo, whereas increases in speed of time were significantly greater for psilocybin in comparison to 2C-B and placebo.

Multidimensional measures of mood, drug effects, and drug liking were taken hourly (see Fig.2, Table 1 and Table S4). Mood as assessed by the POMS (Fig.2a) revealed a greater degree of emotional liability under psilocybin. Psilocybin significantly increased levels across all negative mood subscales (total mood disturbance, tense, anger, fatigue, depression, confusion) compared with placebo. Of these, levels of depression were also significantly greater for psilocybin than 2C-B. Conversely, 2C-B solely elicited significant increases for the tense and confusion subscales in comparison with placebo. 2C-B produced significant increases in markers of positive affect (vigour, elation, friendliness) compared to placebo with the elevations in summary scale positive mood of larger magnitude than under psilocybin. Both compounds generated comparable significant increases on SDRQ scales of current and general drug liking and drug wanting compared to placebo.

**Figure 2.**
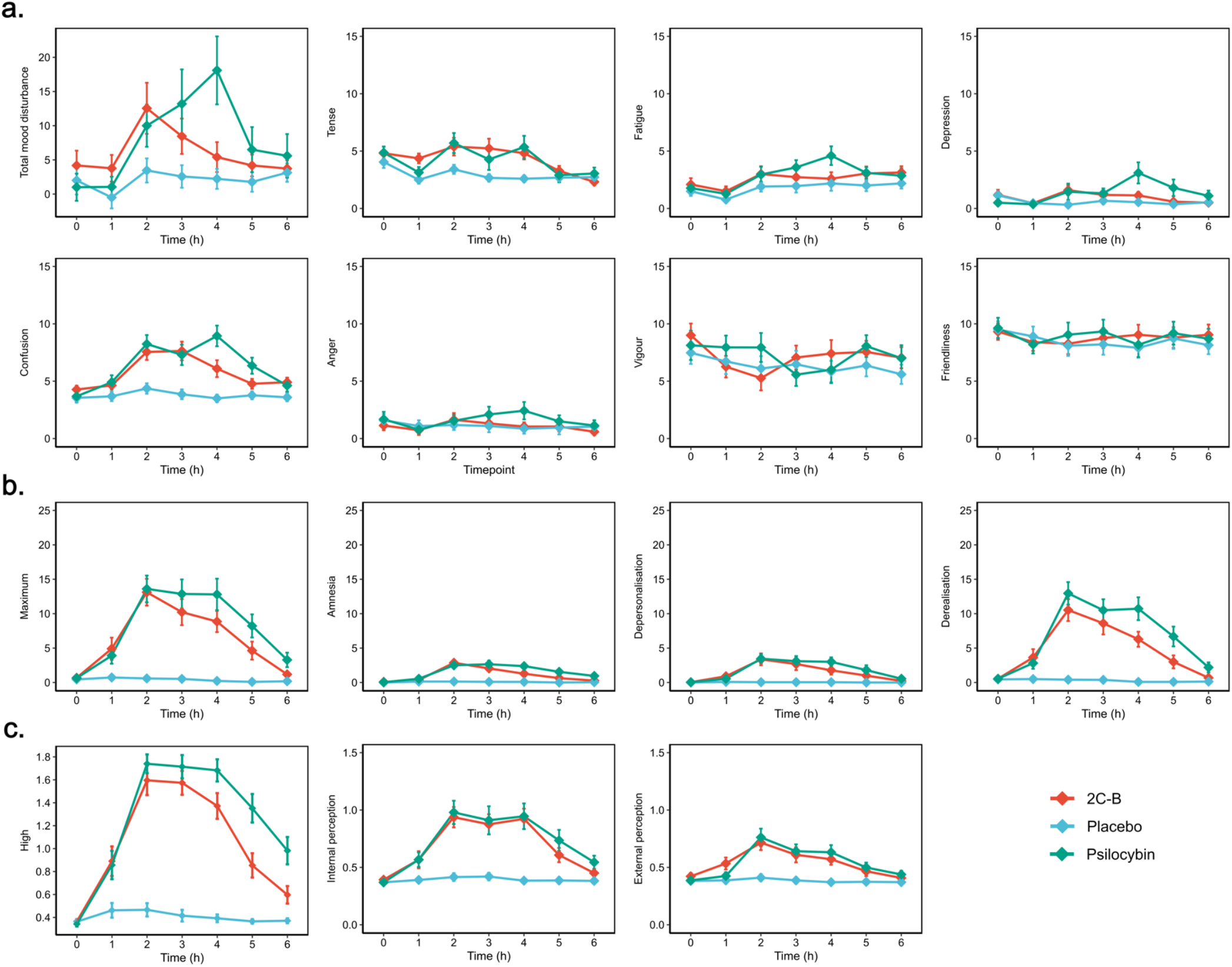
Effect of 2C-B and psilocybin over time on hourly multidimensional measurements of mood, dissociation, and psychedelic effects. Data points are shown as means ± SEM (brackets). Corresponding pairwise statistics for these findings as well as additional subscales are presented in tables 1 and S4. **(a)** Time courses of all primary Profile of Mood States (POMS). **(b)** Time courses of each Clinician-Administered Dissociative States Scale (CADSS) dimension. **(c)** Time courses of the Bowdle Visual Analogue Scale (B-VAS), factors assessing high, changes to bodily and extraneous perception.

Altogether, neither substance significantly differed from one another on any multidimensional measures of acute drug effects. Subjective ratings of dissociation (Fig.2b) and their corresponding subfactors (depersonalisation, derealisation, amnesia) as assessed by the CADSS were significantly greater under both drugs than placebo. Of the subscales, feelings of derealisation exhibited the largest magnitude of change under both drugs. Similarly, real-time scores of internal perception and external perception and high as per the BVAS (Fig.2c) were significantly greater for both substances compared to placebo.

Figure 3 shows participant ratings on three retrospective questionnaires completed approximately 6 hours after drug administration. Statistics are summarised in tables S1 and S4. These results generally show significant increases in measures assessing alterations to waking consciousness after 2C-B and psilocybin. On the 5D-ASC, 2C-B elicited a total alteration in waking consciousness of lesser magnitude in comparison to psilocybin (Fig.3a).

**Figure 3.**
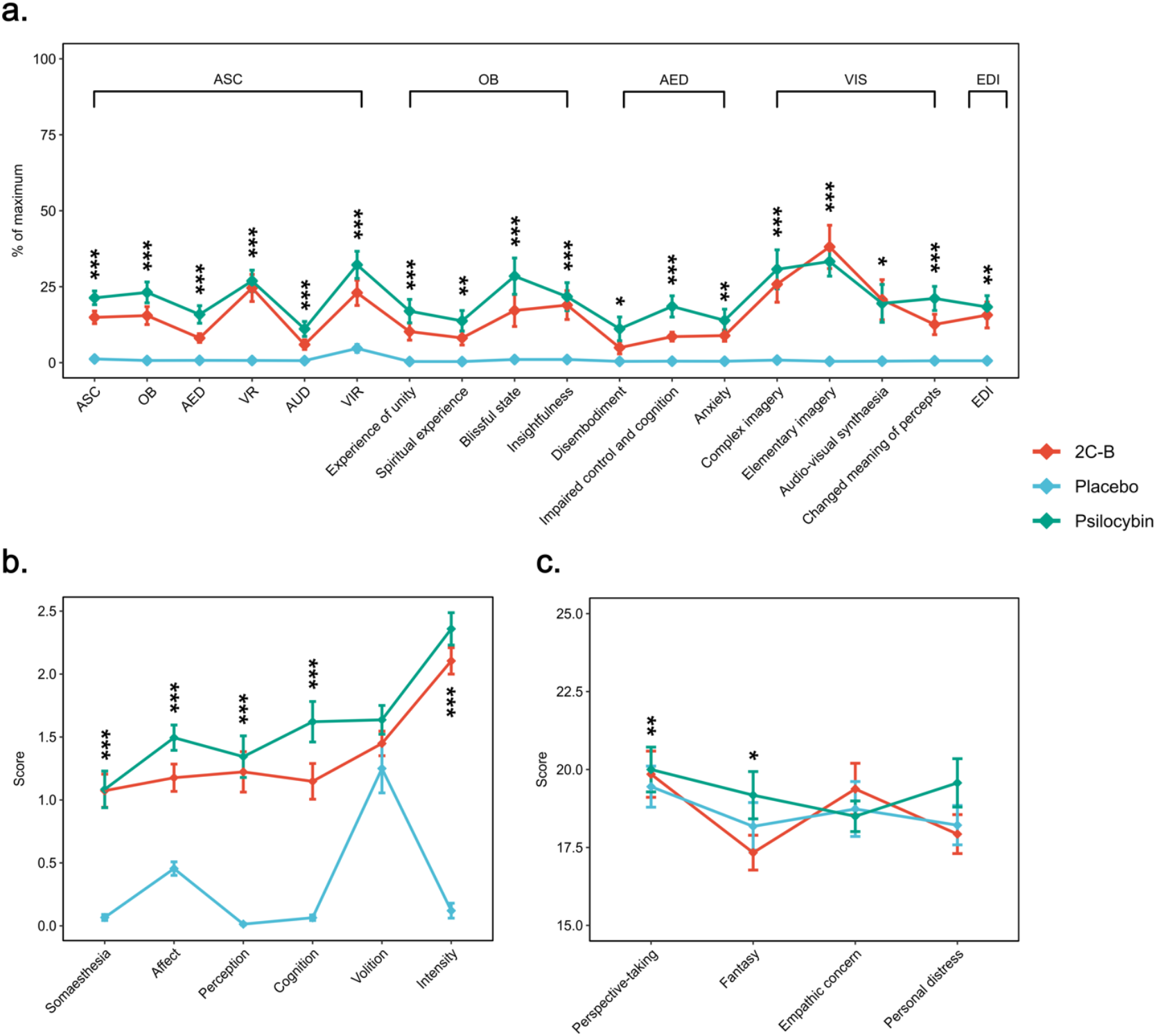
Effect of 2C-B and psilocybin on retrospective measures of changes to waking consciousness and trait empathy. All means ± SEM (brackets) as well as corresponding pairwise statistics are presented in S1 and S4. The presence of significant main drug effect is indicated as follows: \**p* < 0.05, \*\**p* < 0.01, \*\*\**p* < 0.001. **(a)** Combined scores of the 5-Dimensions of Altered States Questionnaire (5D-ASC) alongside the Ego dissolution Inventory (EDI). (**b)** Participant ratings on the Hallucinogen Rating scale (HRS). **(c)** End-of-session differences in trait empathy on the interpersonal reactivity index (IRI).

2C-B generated significantly greater effects for most scales in comparison to placebo, other than for auditory alterations, spiritual experience, and disembodiment. Psilocybin produced significant increases in all dimensions of the 5D-ASC in comparison to placebo. Particularly, psilocybin produced significantly greater increases in the scales oceanic boundlessness (OBE), anxious ego dissolution (AED), auditory alterations (AA), vigilance reduction (VIR), changed meaning of precepts and impaired control and cognition than 2C-B. Both produced significantly greater effects than placebo on the EDI and did not significantly differ from one another (Fig.3a). On the HRS, 2C-B and psilocybin produced significant increases in 4/5 dimensions other than volition (Fig.3b). Notably, psilocybin produced significantly greater increases on the affect and cognition scales than 2C-B. Neither substance significantly differed on the intensity scale of the HRS. Lastly, neither compound elicited any clear immediate changes to trait empathy relative to placebo as indexed by the IRI (Fig.3c).

### Neuropsychological task battery

Altogether, on all cognitive outcome measures, 2C-B and psilocybin comparable effects (see Fig. 4, Table S2, S4). Both produced significant impairments in global cognitive function when compared to placebo as indicated by a reduced number of correct responses and attempts on the DSST. However, no significant effect on general accuracy was identified.

**Figure 4.**
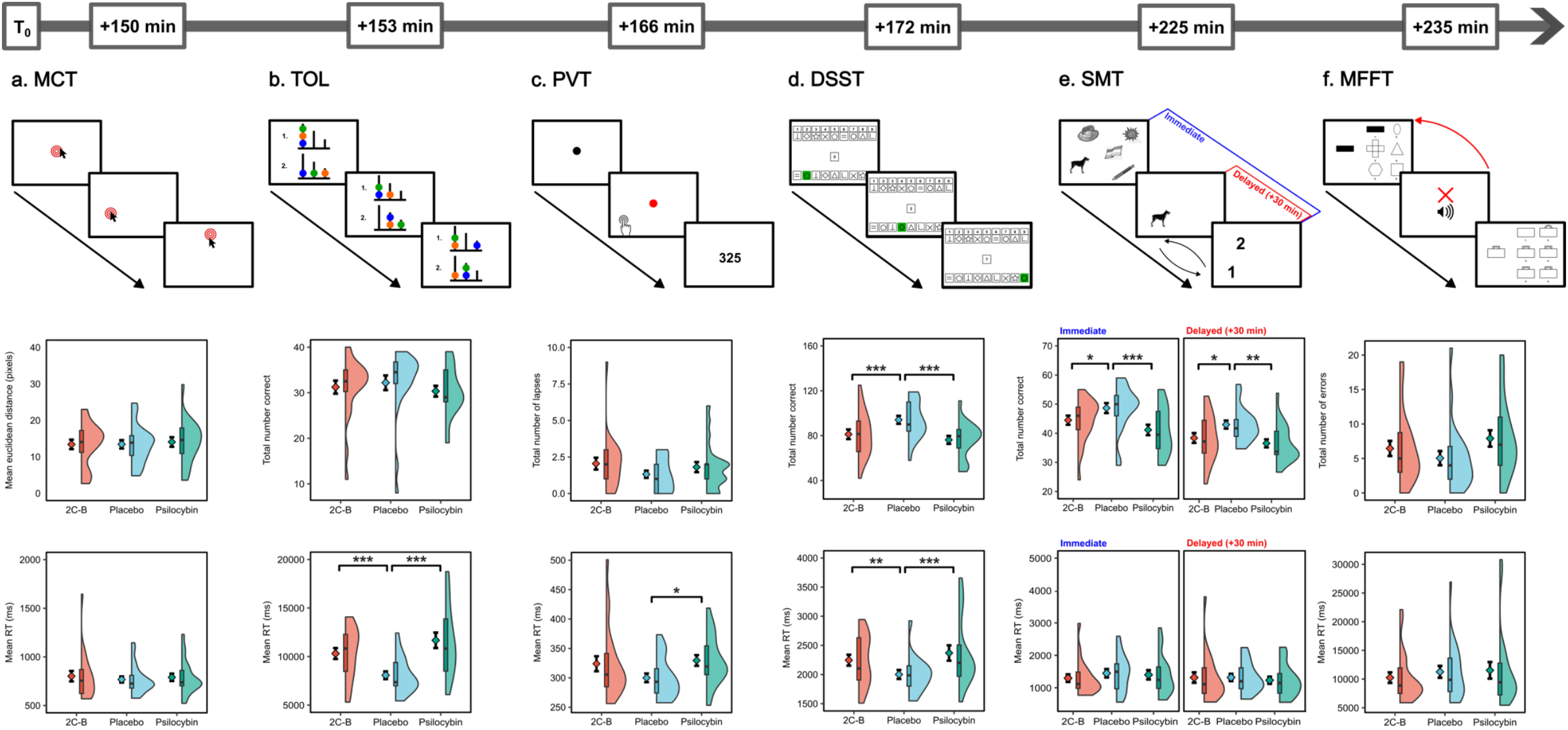
Approximate time course and primary outcomes of task battery. The uppermost panel of each column depicts a schematic of each task as presented to participants with each corresponding time of administration relative to dosing. Lower panels display outcomes as a combined violin plot, boxplot, and mean line (±SEM). In the boxplot, the line dividing the box represents the median, the ends represent the upper 75^th^ / lower 25^th^ percentiles, and the extreme lines represent the full range. Significant pairwise comparisons following a main effect of drug are indicated as follows: \**p* < 0.05, \*\**p* < 0.01, \*\*\**p* < 0.001. **(a)** Motor control task. A target is presented randomly on screen and must be selected as quickly as possible. Participant accuracy (Euclidean distance from target centre) 5) and response times (RTs) are recorded. **(b)** Tower of London. Participants must assess whether the end-arrangement (1>2) can be accomplished in 2-5 steps as quickly as possible. Response times and total correct responses are assessed. **(c)** Psychomotor vigilance task. Subjects must respond to a randomly appearing visual stimulus as quickly as possible. Total attentional lapses (RTs >500 ms) and RTs are registered. **(d)** Digit symbol substitution test. Participants must quickly match novel inputs according to their corresponding digit-symbol combination within 90 seconds, with total correct responses and RTs being the primary outcomes. **(e)** Spatial memory test. Participant must memorise the location of a sequence of 10 targets and individually indicate their correct position (1/2). Subjects are queried 30 minutes later on their location. Total correct responses and RTs are measured. **(f)** Matching familiar figures task. Targets must be matched to one of six possible corresponding alternatives as quickly as possible. Trials are repeated in the presence of an error, with main outcomes being the total number of errors and RTs. All outcomes and corresponding pairwise statistics are presented in tables S2 and S4.

Regarding local facets of cognition, both drugs relative to placebo were associated with significantly diminished scores in immediate and delayed SMT recall. Analyses revealed sensorimotor coordination, mental planning, reflection impulsivity, and empathy (cognitive and emotional) were unaltered under 2C-B and psilocybin, as indicated by the absence of a main effect of drug condition for the MST, PVT, TOL, MFFT and MET (+270 min, see supplementary) respectively. Delayed reaction times in comparison to placebo were selectively present for the DSST and TOL for both drugs as well as the PVT for psilocybin.

### Cardiovascular effects

Acute effects on vital signs over time are shown in Fig. 5a, with peak maximums shown in Table 1. For both compounds, significant pressor effects (systolic and diastolic) were observed in comparison to placebo. Systolic hypertension (>140 mmHg) was observed under 2C-B (n=5), psilocybin (n=8), and placebo (n=2), respectively. Tachycardia (>100 BPM) was observed under 2C-B (n=2) and placebo (n=2) respectively. However, despite a main effect of drug, no clear significant differences in heart rate were observed. Whereas 2C-B produced a significantly greater rate pressure product in comparison to placebo, this did not significantly differ between our active conditions, suggesting an overall similar myocardial output.

**Figure 5.**
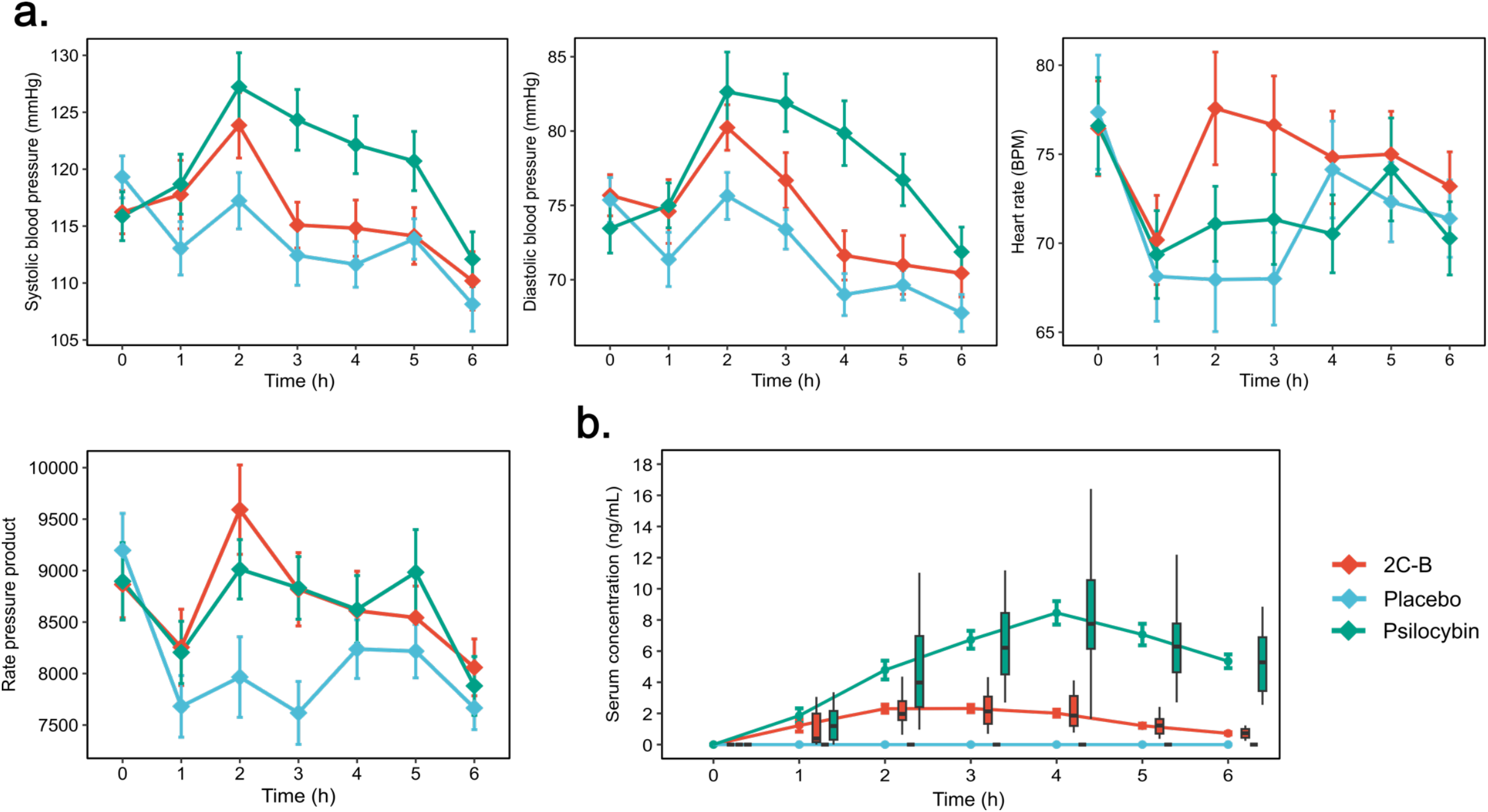
Cardiovascular and pharmacokinetic time-courses of 2C-B and psilocybin. **(a)** Cardiovascular effects over time. Data points are shown as means ± SEM (brackets) with statistical outcomes provided in tables 1 and S4. **(b)** Pharmacokinetic time courses for 2C-B and psilocin. Data points are shown as means ± SEM (brackets) in conjunction with a boxplot for which line dividing the box represents the median, the ends represent the upper 75^th^ / lower 25^th^ percentiles, and the extreme lines represent the full range. **(c)** Effect/concentration hysteresis plots. The pharmacodynamic values are the mean values for any drug effect at each time point in 21 subjects. The time of sampling is noted next to each point (in hours after administration).

### Pharmacokinetics

Concentration time-curves can be found in Fig.5b. The maximum mean values (range, n) for 2C-B and psilocin serum concentrations (C_max_) were 3.31 (1.63–7.58, n=21) and 10.81 (5.26–25.47, n=21) ng/mL, respectively and concordant with applied oral doses^10,35,63^. Maximum serum concentrations peaked for 2C-B on average at 2.43 h (1– 4, n=21) vs 3.71 h (2–5, n=21) for psilocin. Mean AUC_0-6h values_ were 9.43 h*ng/mL (5.03 -16.7, n=21) for 2C-B and 33 h*ng/mL (21.7 - 71.6, n=21) for psilocin. Available mean corresponding half-lives were 1.43 h (1.02 - 2.46, n=17) and 2.24 h (1.44 - 3.81, n=7) for 2C-B and psilocybin respectively. For either drug, exploratory Spearman rank correlations did not reveal significant relationships between C_max_, age, sex nor weight (see supplementary).

### Blinding

Following the completion of each test day, subjects were asked to retrospectively identify the intervention in question. Placebo was correctly identified by 95.5% of subjects, with 2C-B and psilocybin correctly identified by 63.6% of participants. 2C-B was misidentified as psilocybin by 36.4% of subjects. Psilocybin was misclassified as 2C-B for 31.8% of subjects and placebo for 4.5% of subjects (n=1). Data pertaining to blinding identification and decision confidence are shown in in the matrix S3.

## Discussion

The results presented herein provide the first clinical assessment and within-subject comparison of 2C-B and psilocybin across acute markers of subjective, cognitive, and cardiovascular effects in a placebo-controlled fashion. Our results indicate fixed doses of 20 mg 2C-B and 15 mg psilocybin produce broadly similar increases in all acute ratings of peak drug effect intensity, comparable with prior dose-effect studies^9,35^, reflecting a psychotropic equivalence. These findings indicate 2C-B elicits alterations in waking consciousness of lesser experiential depth than psilocybin yet produces comparable cognitive impairment and cardiovascular stimulation.

Globally, 2C-B produced a range of subjective effects consistent with classical psychedelics ^64^. Acute elevations in feelings of dissociation, perceptual change, creativity, and alterations to time perception were observed for both 2C-B and psilocybin. In contrast, nuanced differences in phenomenology were observed across retrospective measurements. Whereas 2C-B was found to elicit significant elevations across most scales outside of auditory alterations, disembodiment, and spiritual experience, the overall intensity of the experience was markedly lesser than that of psilocybin. While exhibiting equivalent alterations to visual (e.g., perception, VR) and bodily perception (ego dissolution, somaesthesia, internal perception), participants under the influence of psilocybin reported a greater propensity for affective qualities of altered self-experience (eg. OBE, AED) as well as impairment (e.g., VIR, impaired control and cognition, cognition). Taken together, these findings support descriptions of 2C-B being non-ego-threatening in nature, lacking the otherwise more serious headspace of classical precursors while imparting a greater emphasis on visual and tactile domains ^19,34^. Consistent with this notion, classification of 2C-B’s 5D-ASC profile in relation to other novel indoleaklylamines has indicated the profoundness of experience as an explanatory factor ^31^. Subtle pharmacodynamic differences between psilocybin and 2C-B may ultimately arise due to differing binding-profiles, given both the autoinhibitory and potentiating interplay of 5-HT_1A_/5-HT_2C_ receptor subtypes on 5-HT_2A_-mediated behavioural responses ^65-68^ or the association of extraneous neuromodulatory systems with feelings of disembodiment and subjective impairment ^69^. Alternatively, it can be suggested the brunt of experiential dissimilarities may instead lie in differing pharmacokinetics such as duration or dosage, given that both 2C-B and psilocybin were unmasked marginally above chance by participants (63.6% of cases). Previous drug-discrimination assays have previously shown a full substitution of most first-generation 2C-X compounds with LSD or DOI ^70^ as well as poor discriminability between psilocybin and LSD in previous comparative studies ^9^.

Whereas classical psychedelics and entactogens partly overlap in regards to their capacity to elicit a positive affective bias ^40^, the former may hold a greater propensity to induce dysphoric reactions due to greater emotional lability. As per prior work^9,35,36,71^, significant elevations in measures pertaining to euphoria (e.g., drug liking, drug high, good drug effect) and friendliness for both 2C-B and psilocybin in comparison to placebo were identified. However, subjects under psilocybin also showed sustained mood disturbance compared to placebo, with significantly larger increases in depression and overall greater emotional range (affect) than 2C-B. Furthermore, compared to placebo, 2C-B did not demonstrate significant elevations in most acute POMS markers of negative affect (anger, fatigue, depression) instead producing elevations in vigour, elation, and positive mood in a similar manner to MDMA ^72^. Seconding this, 2C-B’s effects have been described to resemble a “candy flip,” the concurrent use of MDMA and LSD ^34^. By nature of their action as monoamine reuptake inhibitors, entactogens and psychostimulants are likely to confer lesser susceptibility to negative mood states. For example, noradrenergic effects appear important for their euphoriant nature ^73,74^ while DAT/SERT selectivity has been suggested to distinguish amphetamine-like stimulants from entactogens ^75,76^. In this regard, MDMA and 2C-B have been shown to hold similar DAT/SERT inhibition ratios of 0.08 while stimulants such as amphetamine or methamphetamine show values greater than 10 ^21,77^. Pharmacological challenge studies using duloxetine (a SERT/NET reuptake inhibitor) and reboxetine (a NET reuptake inhibitor) seemingly abolish most mood-enhancing effects of MDMA ^74,78^.

Overall, 2C-B and psilocybin produced a similar profile of cognitive impairment. For the DSST, both compounds yielded reductions in the number of attempted and correct responses, with expectedly longer reaction times. However, neither substance exerted a significant effect on accuracy. Our observations are wholly congruent with work finding psilocybin dose-dependently reduces trial attempts yet spares global accuracy^79^. While we agree with Barrett et al’s 2018 interpretation that the absence of decrements in accuracy is unlikely to be indicative of global executive impairment, we propose parallel reductions in trial attempts and reaction time are another reflection of local impairment. Compensatory psychomotor slowing under high intrinsic cognitive load is likely indicative of deficits in information processing speed rather than a volitional shift in performance strategy^80^ prioritising accuracy over speed. There is also good evidence to suggest disruptions to temporal processing and sensorimotor gating are mediated by 5-HT_2A_ agonism ^81-84^. Deficits in information processing may also consequently explain worsened TOL reaction times in the absence of any main effect on planning. General assessments of executive function such as the DSST are often reliant on a combination of latent cognitive processes such as good motor coordination, spatial memory, sustained attention and response inhibition all of which may be differentially affected according to dose or agent in question ^52^. For example, motor coordination under psilocybin was only previously shown to be affected under high doses (20mg and beyond)^79^. We also observed significant impairments to both short and long-term spatial memory under each compound despite previous evidence of this across serotonergic hallucinogens being mixed ^54,85-87^. Furthermore, any attentional deficits induced by primary 5-HT_2A_ agonists are likely selective: studies employing psilocybin in combination with ketanserin (a 5-HT_2A_/_2C_ antagonist) have specified declines in attentional tracking ability on paradigms requiring distractor inhibition ^85^, suggesting an enhanced salience of distractor stimuli under rather than reduced sustained attentional capacity per se. While disruptions in inhibitory motor responding are seemingly a core feature of serotonergic hallucinogens ^72,84,88^, no effect on MFFT performance was identified, reflecting an unaffected impulsive choice capacity^89^. Similarly, no empathogenic qualities were identified on the MET for either drug, notwithstanding elevations in emotional empathy being cited for similar doses of psilocybin and MDMA ^90^. With both tasks administered at the tail-end of effect intensity, any absence of effect may be a function of time. Going forwards, homing in on clinically relevant domains of cognition will specify our findings further. With alterations in cognitive flexibility being cited as an integral marker of the therapeutic effects of classical psychedelics ^91^, more expansive assessments of executive function such as set-shifting paradigms - may highlight novel differences ^92-94^.

Analogous elevations in blood pressure were identified under 2C-B and psilocybin in comparison to placebo. These were modest in nature and consistent with earlier trials employing classical psychedelics or 2C-B^8,9,35,40^. Magnitudes of change were also appreciably lesser than those observed following the administration of MDMA and related amphetamines ^40,95,96^. Whereas no differences in HR were observed for 15 mg psilocybin as previously described for similar doses ^9,39,97,98^, we did not replicate the prior finding of elevated HR under 2C-B ^35^, likely as a result of differing subject inclusion criteria. These preliminary findings therefore cautiously indicate that the 15 mg psilocybin and 20 mg 2C-B hold similar cardiovascular effects. However, it should be stressed that assessing the full range of 2C-B’s autonomic and endocrine (e.g., cortisol) effects in future phase I trials may be more fruitful for assessing the tolerability of 2C-B.

The kinetics of 2C-B were largely dissociable from those of psilocin. Whereas both compounds demonstrated comparable onset of action times, the effects of 2C-B were shorter than those of psilocybin, mostly abating within 6 hours. This was mirrored by a shorter half -life and smaller AUC for 2C-B in comparison to psilocybin. These findings of low bioavailability may be in agreement with preclinical work suggesting that 2C-B undergoes an extensive first-pass effect after oral administration ^27,99-101^. Body weight had no influence on 2C-B or psilocin serum concentrations, Thus, unlike MDMA^102^ administering 2C-B as a fixed dose may confer equivalent advantages to weight-adjusted dosing similarly to other classical psychedelics ^103^.

Limitations inherent to the study design warrant consideration. Firstly, single doses were used. For a complete categorisation of 2C-B, future trials (eg: NCT05523401) employing an escalating dosing regimen with both MDMA and psilocybin as comparators of interest may provide applicable dosing ranges for healthy subjects. Furthermore, all neurocognitive tasks were performed at fixed timepoints as to accommodate a battery of sufficient breadth. With psychoactive effects being experienced-timeline dependent, differences in actual versus perceived impairment may be reliant on pharmacokinetic differences. However, most tasks were administered at times of approximate peak subjective effects, maintained across the 2 and 4-hour timepoints for both compounds. It is therefore likely tasks administered at these timeframes were not compromised by differences in task timing relative to effect intensity ratings. The present study also had a primary focus on the immediate effects of each compound and by design, did not extend pharmacokinetic sampling beyond 6 hours. Given the subjective effects of psilocybin were not self-resolving by 6h, future studies spanning a full, predefined sampling interval^104^ may more accurately extrapolate population pharmacokinetic parameters such as half-life, clearance or IC_50_. Lastly, testing was performed in a controlled environment, enrolling a homogeneous sample of experienced volunteers. Thus, subjects in different settings, pertaining to different backgrounds may respond in unseen ways to 2C-B or psilocybin. For example, variations have been noted in how individuals perform on the TOL under ayahuasca based on prior use history (where more experienced users show lesser impairment)^105^.

Caveats aside, our findings highlight key considerations for clinical pathfinding. If shown to produce positive mood sequelae akin to classical precursors, novel analogues such as 2C-B which might elicit a mentally “clearer” subjective state, are likely scalable to implement at a clinic, given that acute contact time and dysphoria can be lessened ^106^. For example, evidence of a lessened predisposition to adverse psychological effects may make 2C-B a useful means by which patients apprehensive of macrodoses or at risk of adverse reactions (e.g, high neuroticism) ^107,108^ can be familiarised with a psychedelic-induced subjective experience within a therapeutic model, while retaining resemblances to classical psychedelics in its capacity to induce ego dissolution. Accordingly, dosages of 15-30mg 2C-B have been reportedly employed in both individual and group psycholytic psychotherapy, often adjunct to MDMA or LSD ^109^.

## Conclusion

In summary, these findings give new impetus to the categorisation of 2C-B as a psychedelic with some entactogenic properties. Producing a spectrum of acute subjective, cognitive, and pressor effects compatible with a primary 5-HT_2A_ mode of action, 2C-B produces an effect profile of an intermediary experiential depth. Subsequently assessing it’s acute effects on brain functional organisation, emotional processing and persisting effects will be valuable for understanding its relative harms and clinical utility.

## Author contributions

P.M designed and performed the research, analysed the data, and wrote the first version of the manuscript. N.M designed and performed the research and contributed to the manuscript review. J.R. supported data collection. R.P provided medical supervision and screening. S.T, S.R performed the serum concentration determinations. E.T, K.K designed the research and contributed to the manuscript review. J.G.R designed the research, contributed to the manuscript review and acquired funding.

## Acknowledgments

The authors thank Maria Balaet for providing the MCT, Emma de Brabander, Vitea Medici, Ajna Jansson and Chantal Delaquis for assistance in conducting the study and Cees van Leeuwen for providing medical supervision and screening.

