## Supplementary materials for "Assessment of the acute effects of 2C-B vs psilocybin on subjective experience, mood and cognition"

**Table of Contents:**


- **S1. Methods: Acute subjective effects**
- **S2. Methods: Retrospective subjective effects**
- **S3. Methods: Neuropsychological task battery**
- **S4. Methods: Serum analytics**
- **S5. Methods: Power calculation**
- **S6. Results: CONSORT flowchart**
- **S7. Results: Demographics**
- **S8. Results: Missing data**
- **S9. Results: Duration of effect**
- **S10. Results: MET task performance (+270 min)**
- **S11. Results: Cmax correlations**
- **References**

### S1. Methods: Acute subjective effects

*Subjective effect (VAS)*

Subjective effects were assessed repeatedly using visual analog scales (VASs) at baseline (0h), +0.5, +1, +1.5, +2, +3, +4, +5, +6 hours after drug administration. These scales included the primary VAS items: "*any drug effect*," "*good drug effect*", "*bad drug effec*t", "*drug liking*," "*drug high*", presented as 100-mm horizontal lines (0–100%), marked from "*not at all" on* the left to "*extremely*" on the right [1]. The VAS "*any drug effect*" is an overall effect measure to characterise the overall effect intensity and time course. Prior dose-effect studies have previously demonstrated it to useful marker for pharmacokinetic-pharmacodynamic modeling of psilocybin and LSD's effects [2,3]. Separately, "any drug effect" is interrelated with "drug high", a measure of stimulating effects. The VAS "good drug effect" is an overall measure of positive effects and interrelated with other measures such as "*drug liking*". The VAS "*bad drug effect*" is an overall measure of any negative effects. These items have been demonstrated to sensitive to the acute effects of psychedelics (LSD, psilocybin), entactogens (MDMA, MDA) and psychostimulants (d-amphetamine, mephedrone) as well as 2C-B and 2C-E [1,4-8]. Secondary VAS items comprised "*happy*"," *concentration*"," creative", *"productive*", "*sociable*", *"sense of time*". Items were bidirectional, marked from "*not at all* "*on* the left (0) to "*extremely*" on the right (+100) with "*sense of time*" marked from "*slow*" on the left (0) to "*fast*" on the right (+100). These items have been found to be sufficiently sensitive to enhancing effects at subperceptual doses of LSD [9].

*Profile of Mood States (POMS)*

The POMS consists of 72 adjectives commonly used to describe momentary mood states [10]. Subjects rate from 0 (not at all) to 5 (extremely) the extent to which each adjective describes how they feel at that moment. Item sums produce seven primary factors: Tension, Anger, Fatigue, Depression, Confusion, Vigour, Friendliness and a Total Mood Disturbance score (Tension + Depression + Anger + Fatigue) − Vigour). Derived elements[11] include: Elation, Arousal ((Tense + Vigour) − (Fatigue + Confusion)) and Positive Mood (Elation − Depression). The POMS has previously been used to ascertain the mood-enhancing effects of 2C-B [12] as well as other psychostimulants and entactogens such as 4-FA and MDMA [13,14].

*Clinician-Administered Dissociative States Scale (CADSS)*

The CADSS is an instrument designed to be a standardized measure of present-state dissociative symptomatology [15]. It comprises 19-self report items, ranging in intensity from 0 'not at all' to 4 'extremely present'. The additional seven observer-rated items in the scale, were not employed in the present study. It is comprised of three factors: 1) depersonalisation (feeling detached from oneself), 2) derealisation (feeling detached from one's surroundings) and 3) amnesia (gaps in memory, identity confusion and alteration). Summed together, these subscales form a total dissociative score. The CADSS serves as a rapid assessment of psychedelic effects due to its conceptual overlap with the core loss of subjective self-identity and disembodiment under serotonergic hallucinogens [16-19].

*The Bowdle Visual Analogue Scale (B-VAS)*

The B-VAS is a self-report inventory specifically devised to assess acute psychedelic effects [20-22]. It consists of 13 100-mm VAS from which two composite scales, internal perception (six items) and external perception (five items) are calculated outside of two filler items. The former scale reflects inner feelings not correspondent with reality and the latter scale reflects a misperception of external stimuli or a change in the awareness of the person's surroundings. A third intensity scale - High, is also extrapolated [23].

*Sensitivity to Drug Reinforcement Questionnaire (SDRQ).*

The SDRQ is the adapted form of the Sensitivity to Cannabis Reinforcement Questionnaire, previously used to assess drug desirability [24,25]. This 4-item questionnaire asks participants to rate current and general liking and wanting of the intervention. Subjective valence of liking and wanting is scored from 1 (somewhat) to 5 (extremely).

### S2. Methods: Retrospective subjective effects

*Altered States of Consciousness Rating Scale (5D-ASC)*

The 5D-ASC [26] consists of 94 retrospective 100-mm VAS. The ASC items are grouped into five main dimensions comprising 11 lower-order scales (1) 'oceanic boundlessness' (OB) measures derealization and depersonalization accompanied by changes in affect ranging from heightened mood to euphoria and/or exaltation as well as alterations in the sense of time. The corresponding subfactors include "experience of unity," "spiritual experience," "blissful state," "insightfulness," and "disembodiment." (2) 'anxious ego dissolution' (AED) measures ego disintegration associated with loss of self-control, thought disorder, arousal, and anxiety. The corresponding subfactors comprise "impaired control of cognition", "disembodiment" and "anxiety." (3) 'visionary restructuralisation' (VR) refers to 'elementary hallucinations', 'visual (pseudo-) hallucinations', 'synaesthesia', 'changed meaning of percepts', 'facilitated recollection', and 'facilitated imagination'. It consists of the lower-order scales "complex imagery," "elementary imagery," "audio-visual synesthesia," and "changed meaning of percepts." (4) 'auditory alterations' (AA) refers to acoustic hallucinations and distortions in auditory experiences and (5) the dimension 'reduction of vigilance' (RV) relates to states of drowsiness, reduced alertness, and related impairment of cognitive function. The 5D-ASC is the most widely used psychometric assessment of classical psychedelic and entactogenic effects and has been deployed across a range of altered states of consciousness. Furthermore, acute ratings on the 5D-ASC after administration of psilocybin have been used to predict long-term effects of psychedelic-assisted therapy in patients and healthy volunteers [27,28] while holding good convergence with other clinically used psychometric assessments such as the Mystical Experience Questionnaire [29].

*The Ego-Dissolution Inventory (EDI)*

The Ego-Dissolution Inventory is a self-reported scale used to specifically assess subjective feelings of ego-dissolution/loss of self after drug intake [30]. The questionnaire consists of 8 VAS items (100-mm) which participants have to rate retrospectively, including: I experienced a dissolution of my "self" or ego; I felt at one with the universe; I felt a sense of union with others; I experienced a decrease in my sense of self-importance; I experienced a disintegration of my "self" or ego; I felt far less absorbed by my own issues and concerns; I lost all sense of ego;

All notion of self and identity dissolved away. The EDI has been shown to relate with the underlying effects of 5-$HT_{2A}$ agonists on functional brain organisation [31,32].

*Hallucinogen Rating Scale (HRS)*

The HRS includes 71 items scored 0 (not at all) to 4 (extremely) across six scales: Somaesthesia, reflecting somatic effects including interoceptive, visceral and tactile effects; Affect, sensitive to emotional and affective responses; Volition, indicating the subject's capacity to wilfully interact with his/her 'self' and/or the environment; Cognition, describing impedance in thought processes or content; Perception, measuring visual, auditory, gustatory and olfactory experiences; and finally Intensity, which reflects the strength of the overall experience [33]. The HRS has been extensively used in hallucinogen research employing psilocybin [34,35], DMT [36], ketamine [36,37], Salvia divinorum [38,39], 2C-B [7], MDE (3,4-methylenedioxy-N-ethyl-amphetamine) [40], MDMA [41] and ayahuasca [42,43] as well as psychedelic-assisted psychotherapy studies employing psilocybin [44].

*Interpersonal Reactivity Index (IRI)*

The IRI is a 28-item retrospective inventory using a 5-point scale (1=never; 5=always) which assesses 4 constructs of trait empathy: 'Fantasy' (tendency to imaginatively transpose oneself into fictional situations), 'Perspective-Taking' (capacity to spontaneously adopt the psychological viewpoint of others), 'Empathic Concern', (feelings of sympathy, compassion and concern for others), and 'Personal Distress' (assesses personal distress, unease and anxiety stemming from tense interpersonal settings). The first two scales are a measure of Cognitive Empathy (how well an individual can perceive and understand the emotions of another), the latter two ('Empathetic Concern and 'Personal Distress') assess Emotional Empathy, reflecting emotional contagion [45]. The IRI was employed as a confound to assess whether any acute state-dependent empathy-enhancing effects of 2C-B or psilocybin (as assessed by the multifaceted empathy test (MET) arise independently from fluctuations in a subject's trait-level empathy [46].

**S3. Methods: Neuropsychological task battery**

Impairments in cognitive domains pertinent to the acute effects of psychedelics, entactogens and psychostimulants were assessed using a computerised task battery delivered on a dedicated testing laptop at fixed timepoints throughout the test day.

*The Motor Control Task (MCT)*

The MCT is an analogue of the CANTAB Motor Screen Task [47] and provides a general assessment of whether sensorimotor deficits will limit the collection of valid data from the participant. Coloured targets are presented in different locations on the screen, one at a time. The participant must select the target on the screen as quickly and accurately as possible. Outcome measures assessed include a participant's reaction time and the accuracy of pointing (selecting the cross), the latter defined as the mean Euclidean distance in pixels from the target centre.

*Tower of London (TOL)*

The TOL serves as a measure of executive functioning, specifically dimensions pertaining to mental planning and decision making [48,49]. Task difficulty is dictated by elements such as for example the minimum number of moves required to complete the goal, goal hierarchy (i.e., ambiguity of the sequence of final moves derived from the configuration of the goal state). The present version consists of computer-generated images of begin- and end-arrangements of three coloured balls. Every individual movement of the ball is counted as one step. Participants decide as quickly as possible, whether the end-arrangement can be accomplished in 2, 3, 4 or 5 steps from the begin arrangement by pushing the corresponding coded button. Reaction times and total correct responses are the main dependent variables. Separate counterbalanced versions comprising unique problem sequences were provided for each test day. The TOL has been previously employed in acute studies administering ayahuasca [50].

*Psychomotor Vigilance Task (PVT)*

The PVT assesses reaction times in response to a visual stimulus as a measure of sustained attention [51]. The visual stimulus is a red circle presented at random intervals. Participants must press a button as quickly as possible upon its onset. Outcome measures were mean reaction time (milliseconds) and the number of attentional lapses. An attention lapse is the failure to react, or a reaction time exceeding 500 ms. Typically employed in circumstances of diminished arousal, such as sleep deprivation, the PVT has also been used to assess the stimulating effects of MDMA and d-amphetamine [52-54].

*Digit Symbol Substitution Test (DSST)*

The DSST is a computerised version of the original paper and pencil test taken from the Wechsler Adult Intelligence Scale [55]. The participant is shown an encoding scheme consisting of a row of squares at the top of the screen, wherein nine digits are randomly associated with particular symbols. The same symbols are presented in a fixed sequence at the bottom of the screen as a row of separate response buttons. The randomisation procedure is chosen such that symbols never appear at the same ordinal position within both rows. The encoding scheme and the response buttons remain visible while the participant is shown successive presentations of a single digit at the centre of the screen. The goal is to match each digit with a symbol from the encoding list and click the corresponding response button. The number of digits correctly encoded within 3 minutes is the primary outcome. Secondary outcomes include number of attempts, percentage accuracy (total correct/total attempts) and reaction time. Unique counterbalanced versions of the task were deployed on each test day, differing in symbol types and ordering. The DSST has been employed in assessments of cognitive impairment under psilocybin, MDMA, dissociatives (ketamine, dextromethorphan), psychostimulants (mephedrone,4-FA,d-amphetamine) [13,56-60].

*Spatial Memory Test (SMT)*

The SMT consists of an immediate and a delayed recognition phase, serving to evaluate visuospatial memory and reasoning in psychopharmacological drug trials [61,62]. The immediate recall phase is composed of six trials in which ten black-and-white pictures (total 60 pictures) are subsequently presented on a computer screen at different locations for a duration of 2 s and inter-stimulus interval of 1 s. After every trial, the pictures reappear one by one in the middle of the screen for 2 s followed by the presentation of a “1” and a “2” in different locations. The participant must indicate whether each picture corresponded with either location 1 or 2. The delayed recognition phase is completed after a 30 min delay in which the participant must indicate the correct picture location.

*Matching Familiar Figures Test (MFFT)*

The Matching Familiar Figures test (MFFT) is an information sampling task assessing reflection-impulsivity [63-65]. This task was developed to assess the processes involved in the gathering and evaluation of perceptual information required to make a response. Subjects are simultaneously presented with a target figure on the left-hand side of the screen alongside six alternative figures on the right-hand side which differ in one or more details, all except one. Subjects must identify and select the matching alternative as quickly as possible by pressing the corresponding number on the keypad. If the initial choice is incorrect, a beep is played, and participants are required to repeat the trial. The task comprises 20 test trials preceded by 2 practice trials. Primary outcomes include mean latency to first response and total number of errors. Additional outcomes include: an Impulsivity score (I-score) serving as a composite score of impulsivity and an Efficiency score (E-score) which serves as an index of the balance between “fast and accurate” vs “slow and inaccurate” responding [66]. Counterbalanced versions of the task were employed for each dosing day, comprising novel figures. The MFFT has been previously used to assess impulsive action under MDMA and psychostimulants [14,25,67].

*Multifaceted Empathy Test (MET)*

The MET consists of 40 pictures of people conveying a complex emotional state which was positive in 50% of the pictures and negative in the other half. To assess cognitive empathy, participants had to select, out of 4 words, the emotion word which matched the picture. To assess emotional empathy (EE), participants had to rate on a scale from 1-9 ‘how aroused this picture made them feel’ (Implicit EE) and ‘how concerned they were for the person’ (Explicit EE). Primary outcomes comprise the number of correct classified pictures and the IEE/ EEE ratings per valence [68]. The MET has been previously used to identify empathogenic qualities under entactogens and classical psychedelics [69].

**S3. Methods: Serum analytics**

All samples were centrifuged, and serum was stored frozen at -20°C until analysis.

Analysis of 2C-B. Serum (200 µl) was extracted with 1 ml of ethyl acetate/methyl tertiary-butyl ether (80:20, v/v) after addition of 0.2 ml phosphate buffer pH 9 and 50 µl of internal standard (acetonitrile containing 0.5 ng of 2C-B-d6 from Toronto Research Chemicals, Toronto, Canada). Analysis of psilocin in serum was performed according to Martin, et al 2013.[70]. Serum (200 µl) was extracted with 1 ml of ethyl acetate after addition of phosphate buffer pH 9, 20 ng psilocin-d 10 and 10 µl of 0.1 M ascorbic acid for stabilization.

For both analysis streams, the organic phase was evaporated and reconstituted with 100 µl of 0.1 % formic acid/acetonitrile (80:20, v/v). The analysis of (5 µl 2C-B, 2 µl psilocin) was performed on an Agilent (Waldbronn, Germany) LC-MS/MS system consisting of a 1290 Infinity Liquid Chromatograph coupled via JetStream Electrospray Interface (ESI) to a G6460A Triple Quadrupole Mass Spectrometer. Analytes were separated on a Kinetex® 2.6 µm XB-C18 100 Å LC column (100 x 2.1 mm) plus corresponding guard column from Phenomenex (Aschaffenburg, Germany) at 30 °C. Gradient elution at a flow rate of 0.5 ml/min using 0.01% formic acid containing 5 mM ammonium formate (A) and acetonitrile containing 0.1 % formic acid (B) started with 5 % B, increased to 95 % B during 4 min and was held for 2 min. Source parameters were: gas temperature 300 °C, gas flow 11 l/min, nebulizer 45 psi, sheath gas temperature 400 °C, sheath gas flow 12 l/min and capillary voltage 3500 V. Detection was performed in the multiple reaction monitoring mode (m/z, collision energy in parentheses, quantifier underlined): 2C-B-d6: 266→249.0 (8V); 2C-B 260→243 (8V); 228 (20V); psilocin-d10: 215®66 (12), psilocine 205®58 (12); 205®160 (16). Seven calibration standards were prepared in the range 0.2 – 50 ng 2C-B and five calibration standards in the range 1 – 100 ng/ml for psilocin per ml human serum and were analysed with the samples. Calibrations were linear (regression coefficients >0.99).

#### S4. Methods: Power calculation

The primary study outcome of this trial (CCMO register: NL73539.068.20) was defined as the number of correct responses on the DSST as a measure of global impairment, due to its correspondence with the remaining neurocognitive test battery. A G*Power calculation based on a repeated mixed effect model study was determined by inputting 3 levels of treatment: 2 active conditions ( 2C-B, psilocybin) and 1 inactive condition (bitter lemon placebo), revealing that given a significance level of $p$ =0.05, a minimal expected effect size of 0.33 and power set at 90%, a minimum of 12 participants were needed to significantly detect differences in task performance between our active and placebo treatments.

#### S5. Methods: Screening procedure

Preliminary eligibility was determined via online prescreening (Qualtrics XM) assessing prior use history, frequency, and location. If eligible, subjects were invited for an online debriefing of the study procedures and measures. After obtaining written informed consent, subjects were sent digital evaluations of their health history and drug use history. If consistent with the study inclusion criteria, subjects were invited for a more in-depth in-person evaluation. The following was performed; thorough evaluation of the patient's physical and mental health, vital signs, weight, and electrocardiogram were recorded, and a physical examination was conducted. Laboratory tests performed included: urine toxicology for street drugs, urinalysis, serum chemistry, hematology, and liver function. Participants were told that while they may not benefit from study participation, their participation may lead to knowledge that may help others.

### S6. Results: CONSORT flowchart

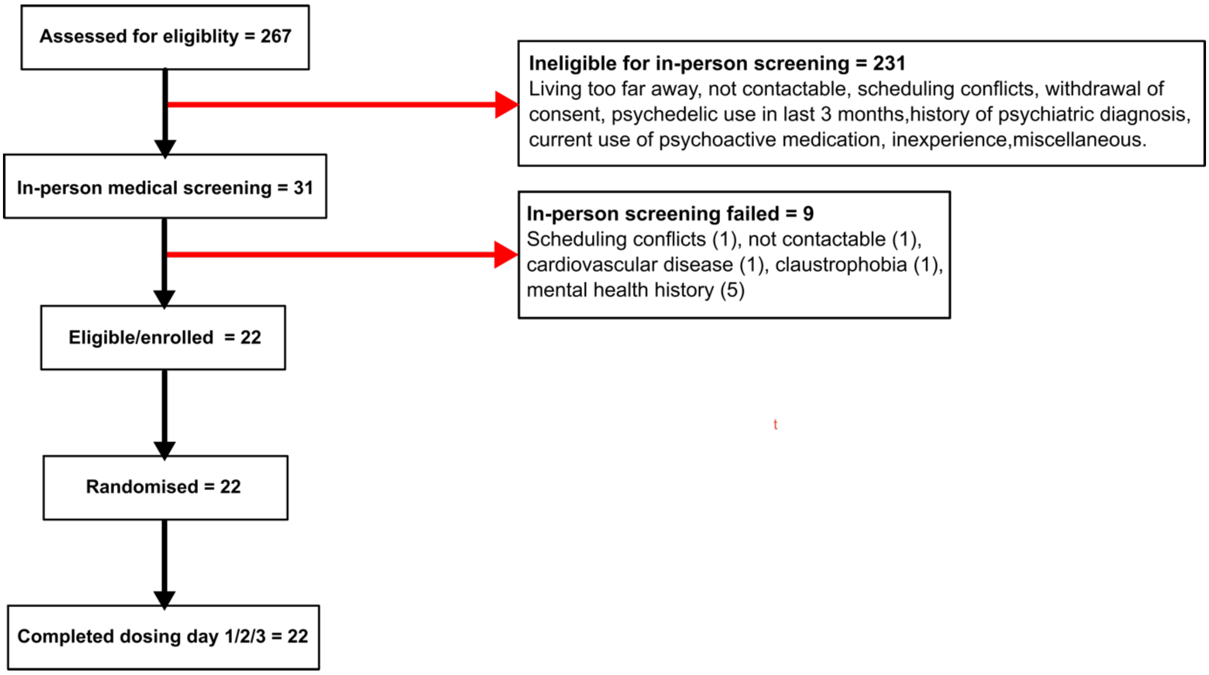


### S7. Results: Demographics

| ***Variable (n = 22)*** | ***Mean (SD)*** |
|---|---|
| Sex (male/female), n | 11/11 |
| Age, years | 25 (4.00) |
| Weight, kg | 69.94 (10.07) |
| History of psychedelic use, years | 4 (2.00) |
| Lifetime psychedelic use, number of occasions | 24 (41.00) |
| Cannabis consumption, per month | 2.12 (1.94) |
| Alcohol consumption, glasses per week | 2.80 (3.70) |
| Caffeine consumption, glasses per week | 12.55 (9.64) |
| Nicotine consumption, cigarettes per week | 2.73 (9.75) |

#### S8. Results: Missing data

| *Outcomes* | | *N of total responses across dosing visits (n = 66)* |
|---|---|---|
| Cognition | MST | 2 |
| | TOL | 1 |
| | PVT | 1 |
| | DSST | 3 |
| | SMT - immediate | 3 |
| | SMT – delayed | 5 |
| | MFFT | 1 |
| | MET | 1 |
| Pharmacokinetics | | |
| | No available data | 2 |
| | Missing timepoints | 3 |

#### S9. Results: Duration of effect

We extrapolated the onset, temporal maximum, and duration of subjective effects for 2C-B and psilocybin for each subject by on/off cut-off of 10% of the maximum individual response for the VAS “Any drug effect”. Effect onsets were 1.20 (SD: ±0.95 and 1.16 (±0.70) respectively with absolute Tmaxs arising at 2.55 (SD: ±1.02) and 3.32 (SD: ±1.24) for 2C-B and psilocybin.

### S10. Results: MET task performance (+270 min)

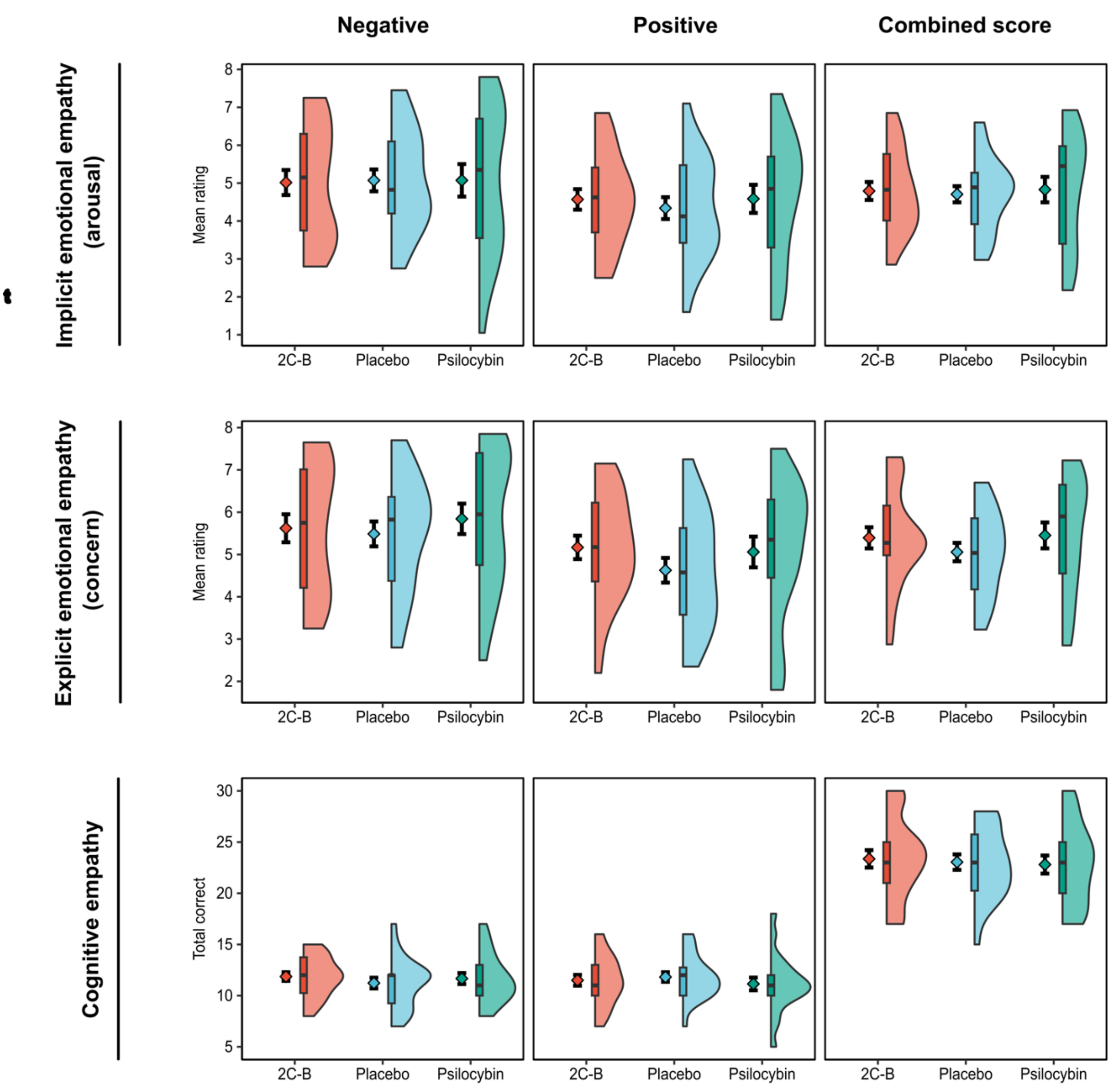


*Multifaceted empathy test (MET) performance at +270 minutes.* Scores for the MET dimensions of emotional empathy (implicit & explicit) and cognitive empathy are plotted on combined split-half violin plots. The plot is a combination of a probability density plot, boxplot, and mean line (± standard error of mean). In the boxplot, the line dividing the box represents the median of the data, the ends represent the upper 75th / lower 25th percentiles, and the extreme lines represent the highest and lowest values excluding outliers. No significant effect of drug was identified for any outcome measure of the MET (see supplementary table S3).

### S11. Results: Cmax correlations

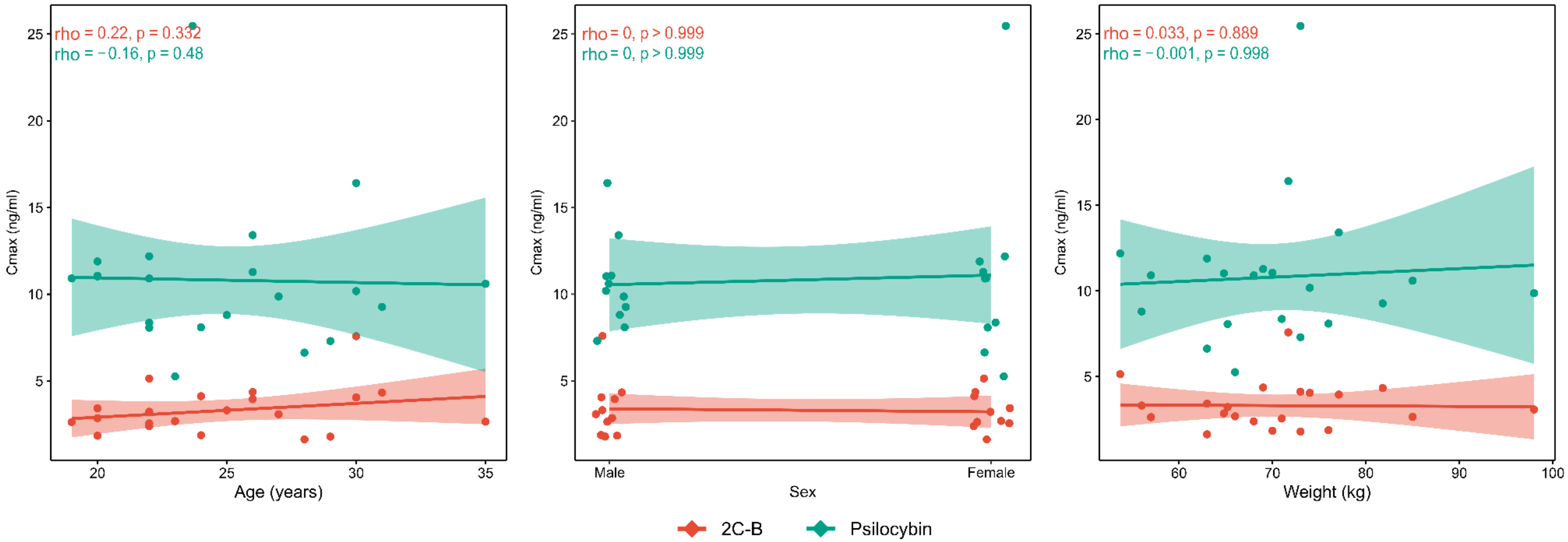


*Spearman rank $C_{max}$ correlation scatterplots.* Raw datapoints are displayed per drug, alongside linear trend lines. Spearman rank correlations were performed for age, sex and weight versus serum drug concentration $C_{max}$. Sex associations were assessed using a point biserial spearman rank correlation approach in order to account for the binomial distributions of sex. Line shadings reflect correlation confidence intervals. Spearman rho's and corresponding p-values are made available for each plot.  No significant associations were identified.
